# Near-chromosomal *de novo* assembly of Bengal tiger genome reveals genetic hallmarks of apex-predation

**DOI:** 10.1101/2022.05.14.491975

**Authors:** Harsh Shukla, Kushal Suryamohan, Anubhab Khan, Krishna Mohan, Rajadurai C. Perumal, Oommen K. Mathew, Ramesh Menon, Mandumpala Davis Dixon, Megha Muraleedharan, Boney Kuriakose, Saju Michael, Sajesh P. Krishnankutty, Arun Zachariah, Somasekar Seshagiri, Uma Ramakrishnan

## Abstract

The tiger, a poster child for conservation, remains an endangered apex predator. Continued survival and recovery will require a comprehensive understanding of their genetic diversity and the use of such information for population management. A high-quality tiger genome assembly will be an important tool for conservation genetics, especially for the Indian tiger, the most abundant subspecies in the wild. Here, we present high-quality near-chromosomal genome assemblies of a female and a male wild Indian tiger (*Panthera tigris tigris*). Our assemblies had a scaffold N50 of >140□Mb, with 19□scaffolds, corresponding to the 19 numbered chromosomes, containing 95% of the genome. Our assemblies also enabled detection of longer stretches of runs of homozygosity compared to previous assemblies which will improve estimates of genomic inbreeding. Comprehensive genome annotation identified 26,068 protein-coding genes, including several gene families involved in key morphological features such as the teeth, claws, vision, olfaction, taste and body stripes. We also identified 301 microRNAs, 365 small nucleolar RNAs, 632 tRNAs and other noncoding RNA elements, several of which are predicted to regulate key biological pathways that likely contribute to tiger’s apex predatory traits. We identify signatures of positive selection in the tiger genome that are consistent with the *Panthera* lineage. Our high-quality genome will enable use of non-invasive samples for comprehensive assessment of genetic diversity, thus supporting effective conservation and management of wild tiger populations.

## Introduction

Tigers are among the most iconic and recognizable species in the world. Despite being apex predators, they are among the most endangered animals with an estimated global population of 3,900 tigers remaining in the wild compared to over 100,000 at the turn of the 20^th^ century (https://www.worldwildlife.org/species/tiger). Historically, wild tigers roamed large swaths of the planet that included a range spanning present-day Armenia in Eastern Asia to Indonesia in Southeast Asia and from the Russian Far East to the Southern tip of India [1]. Anthropogenic activities such as hunting, urbanization, expansion of agriculture and deforestation, leading to both loss of habitat and prey have resulted in a >95% decline in the wild tiger populations [2].

The Bengal tiger or the Indian tiger, *Panthera tigris tigris*, is a subspecies native to the Indian subcontinent and is highly endangered. It is estimated that over 50,000 tigers inhabited the unbroken forests of India [3]. Much like global tiger population decline, Bengal tiger population in India also began to decline more than a century ago, leaving fewer than 2,000 tigers in the wild by 1970. In 1973, India declared the tiger its national animal and set up ‘Project Tiger’ to conserve this majestic animal [4]. With over 50 tiger reserves, the project has given the tiger a chance for continued survival. Today there are ~2,900 wild tigers in these reserves in India, accounting for ~60% of the global wild tiger population [5]. The Bengal tiger population is estimated to be genetically the most diverse and hence represents the best gene pool reservoir important for conservation [1, 6, 7]. Since genetic diversity in a population takes times to build up, for Bengal tigers, small and fragmented reserves averaging <1,500 square kilometers in India may lead to isolated population groups susceptible to inbreeding depression and genetic, demographic and environmental stochasticity there by posing a conservation challenge [1, 6]. In this context, continued monitoring of the genetic diversity of these populations will be important. Obtaining high quality samples from the animals in the wild to conduct genetic studies is not practical. A high-quality reference genome will allow the use of DNA from non-invasive samples scat samples to study, monitor and manage wild populations [8].

The evolution of genomics techniques/technologies over the last decade [9, 10] enable high-quality *de novo* genome assemblies of endangered and threatened species [11–15]. These include short- and long-read sequencing, optical mapping and chromosome conformation capture sequencing technologies that together allow for the generation of near-chromosomal genome assembly.

The Bengal tiger is a diploid organism with a chromosome number 2n = 38, which includes 16 pairs of metacentric/submetacentric autosomes, two pairs of acrocentric autosomes, a meta-centric X, and Y sex chromosome. To date, none of the published tiger genomes are near-chromosomal assemblies. The Amur tiger (*Panthera tigris altaica*) genome assembly, generated using short-read data was highly fragmented [16]. Recently, a genome assembly for a captive-wild-caught Malayan tiger (*Panthera tigris jacksoni*) using 10x linked-reads has been reported [6]. Additionally, although several tiger resequencing studies have produced additional sequence data [8, 17], none have resulted in a contiguous high-quality reference genome.

Here, we report near-chromosomal *de novo* reference genome assemblies of a male and a female Bengal tiger from the wild. The female animal is a famous Bengal tigress named Machali from the Ranthambore National Park, India, crowned the Queen Mother of Tigers and is the most widely photographed tiger in the world (https://en.wikipedia.org/wiki/Machali_(tigress)). Machali, a reproductively successful founder, gave birth to seven females and four males between 1999 to 2006.

We have performed a comprehensive annotation of our high-quality genomes and identified several genes that contribute to key traits that make the tiger an apex predator. This included genes involved in skin patterning, tooth development, vision, endurance and olfaction. We also show that the contiguous genomes generated here enable identification of longer stretches of runs of homozygosity (ROH) compared to previousl reported assemblies, a key measure of inbreeding. Further, we used the reference quality gene annotation models to analyze signals of positive selection and evolution of big cat-specific traits.

## Results

### Genome sequencing and chromosome-level assembly

We generated high-quality genome assembly using long-read and short-read sequencing data, chromatin conformation (Hi-C) sequence and/or optical mapping (Bionano) data (**Supplementary Table 1a-b**, materials and methods) [18, 19]. The Machali (MC) draft assembly generated using long-read sequence data contained 2,766 contigs, spanning 2.4 Gb and had a contig N50 of 4.40 Mb (**Table 1a**; see materials and methods). Following error correction using PacBio and Illumina data, we obtained the PanTigT.MC.v1 assembly that served as input for Hi-C data scaffolding [20]. This resulted in a near-chromosomal assembly, PanTigT.MC.v2 containing 1,052 scaffolds and a scaffold N50 of 145.47 Mb. This represented a ~50x improvement in genome contiguity (see materials and methods). The BUSCO (v5.3.2) completeness score for this assembly for detection of conserved carnivora single-copy orthologs (**Table 2**) was 95.1% [21, 22].

**Table 1:** Genome assembly statistics for (a) MC and (b) SI genomes.

| Assembly | Size<br>(Gb) | # of<br>contigs | # of<br>scaffolds | Contig/scaffold<br>N50 (Mb) | Longest<br>scaffold (Mb) |
| --- | --- | --- | --- | --- | --- |
| PanTigT.MC.v1 (LR + SR) | 2.41 | 2,766 | - | 4.40 | 21.20 |
| PanTigT.MC.v2 (v1 + Hi-C) | 2.41 | - | 1,052 | 145.47 | 237.64 |

**b)**
| Assembly | Size<br>(Gb) | # of<br>contigs | # of<br>scaffolds | Contig/scaffold<br>N50 (Mb) | Longest<br>scaffold (Mb) |
| --- | --- | --- | --- | --- | --- |
| PanTigT.SI.v1 (LR + SR) | 2.38 | 845 | - | 29 | 131.97 |
| PanTigT.SI.v2 (v1 + Chicago) | 2.53 | - | 6,889 | 2.10 | 29.57 |
| PanTigT.SI_BNG (BNG) | 2.63 | - | 182 | 145.93 | - |
| PanTigT.SI.v3 (v2 + BNG) | 2.53 | - | 3,808 | 147.30 | 238.82 |
| ONT+Chicago+Bionano+Hi-C | 2.53 | - | 3,808 | 2.41 | 238.82 |

**Table 2:** Genome assembly statistics.

| Assembly | Bengal Tiger<br>(PanTigT.MC.v3) | Bengal Tiger<br>(PanTigT.SI.v3) |
| --- | --- | --- |
| Total sequence length (Gb) | 2.41 | 2.53 |
| Contig N50 (Mb) | 4.40 | 2.41 |
| Number of scaffolds | 1,052 | 3,808 |
| Number of scaffolds > 10 Mb<br>(% of assembly) | 19 (94.1) | 19 (93.3) |
| Largest scaffold (Mb) | 237.6 | 238.8 |
| Scaffold N50 (Mb) | 145.47 | 147.0 |
| Number of gaps | 1700 | 5708 |
| BUSCO (%) | 95.1 | 91.3 |

Similarly, the initial draft assembly of the Southern Indian (SI) animal obtained using long-read data consisted of 845 contigs with a contig N50 of 29 Mb (**Table 1b**, **Supplementary Table 1a-b**; see materials and methods). After polishing with Illumina short-read sequencing, we used the resulting assembly, PanTigT.SI.v1, as input for scaffolding with Chicago chromatin interaction mapping data and ordered and oriented the contigs, corrected mis-joined and merged overlaps. Integration with Chicago data led to a resolved assembly N50 of 2.3 Mb (PanTigT.SI.v2; **Table 1b**). This is consistent with the finding that Chicago data captures short range chromatin interactions and corrects for misjoins in assemblies obtained using long-read sequencing data alone [20]. Integration of this assembly with *de novo* assembled Bionano optical map data (PanTigT.SI_BNG) resulted in an assembly, PanTigT.SI.v3, containing 3,808 scaffolds and a scaffold N50 of 141.0 Mb. The BUSCO (v5.3.2) completeness score (carnivora) for this assembly was 91.3% (**Table 1b and 2**). Interestingly, further incorporation of Hi-C data did not result in an improvement of the SI genome contiguity (see materials and methods).

To assess for any ordering/orientation errors in the PanTigT.MC.v2 assembly after Hi-C scaffolding, we used the chromosome-level reference genome of domestic cat (*Felis catus*; Genbank: GCA_013340865.1, *Felis catus 9.0*) [23] as well as the PanTigT.SI.v3 assembly to delineate the errors from actual structural changes (see materials and methods). Given the highly conserved karyotype among Felidae species [24], we used the domestic cat reference assembly to merge scaffolds into chromosomes and assign chromosomes based on synteny giving rise to the final near-chromosomal assemblies, PanTigT.MC.v3 and PanTigT.SI.v3 (**Figure 1a-b**). The largest scaffold from PanTigT.MC.v3 and PanTigT.SI.v3 spanned over 230 Mb. Further, using the male PanTigT.SI.v3 assembly, we were also able to identify 3 Y-chromosome-linked scaffolds that spanned ~3 Mb (**Figure 1c**; see materials and methods). Overall, both genome assemblies were highly contiguous, with >93% of the genome contained in the 19 near-chromosome-level scaffolds and were syntenic with each other (**Supplementary Figure 1**). Whole genome alignment of the two genome assemblies using the MC genome as the reference revealed 2,196,239 SNPs (~1.05 SNPs/kb) in the SI genome (see materials and methods). Also, consistent with the evolutionary history, both genomes were highly collinear with the domestic cat reference genome (**Figure 1d, Table 2**).

**Figure 1:**
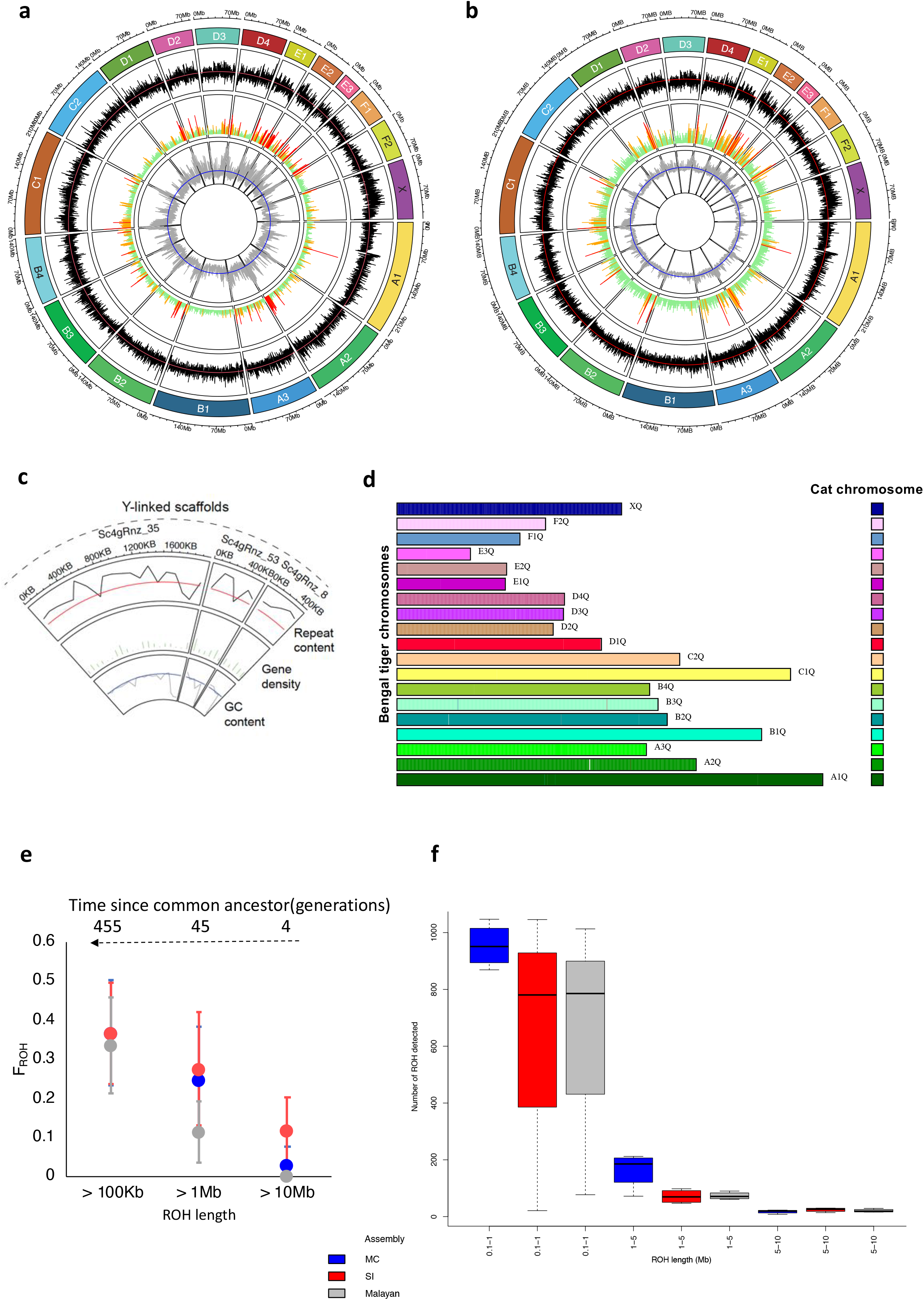
Bengal Tiger Genome assembly. (**a-b**) Circos plot representing the near-chromosome-level assembly of female (MC) **(a)** and male (SI) (**b**) Bengal tiger genomes. The two outermost tracks represent the chromosome length (Mb) and ID. Repeat content per 100 kb window per chromosome (red line represents the mean genome-wide mean repeat content ~36 %) is shown in black. Gene density (green lines indicate 1-5 genes per 100 kb window per chromosome, orange 5-10, and red > 10 genes) is displayed next. The innermost track represents GC% per 100 kb window per chromosome (blue line represents mean GC% - ~41%). (**c**) Y-linked scaffolds from the male tiger genome assembly. Circos plot track as in (a-b). (**d**) Chromosome painting showing synteny between domestic cat genome (FelCat9.0; Genbank: GCA_000181335.4) and the female Bengal tiger (MC) scaffolds. (**e**) Genomic inbreeding coefficients (F_ROH_) derived from runs of homozygosity (ROH) of different lengths (>100 Kb, >1 Mb and >10 Mb) and (**f**) box plot of binned distribution of ROH lengths in the four zoo-bred tigers as derived when using the MC, SI and Malayan tiger genomes as the reference genome.

Comparison of the MC and SI genome assemblies to other published Felidae genomes showed that the overall contiguity was better than the domestic cat reference genome. For comparison, the scaffold N50 of the Bengal tiger genomes were ~1.7× (83.8 versus 145-147 Mb) longer than that of the domestic cat genome (Felis_catus_9.0, Genbank: GCA_013340865.1), a gold standard feline genome. Unlike the tiger genome assembled *de novo* in this study, the domestic cat genome was assembled using physical mapping and sequencing. Also, it was improved iteratively since its publication in 2007 [25]. A comparison of our genome assemblies to that of the recently published Malayan tiger genome (*Panthera tigris jacksoni*) showed that they were ~7x (21.2 versus 145-147 Mb) better as assessed by scaffold N50 [6, 16](MalTig1.0, Genbank: GCA_019348595.1) (**Table 3**). Also, a comparison of our genome assemblies to that of the Amur tiger genome showed that they were ~16×□more contiguous (8.8 versus 145-147□Mb scaffold N50, respectively) [6, 16] (PanTig1.0, Genbank: GCA_000464555.1) (**Table 3**). Although, genomes of other big cats such as jaguar and leopard from the *Panthera* genus have been published [26, 27], none are near chromosomal reference assemblies (**Table 3**). Given our assembly included a male Bengal tiger, we were able to identify Y chromosome-associated scaffolds that provides an important a reference for Y-haplotyping of tiger populations.

**Table 3:**
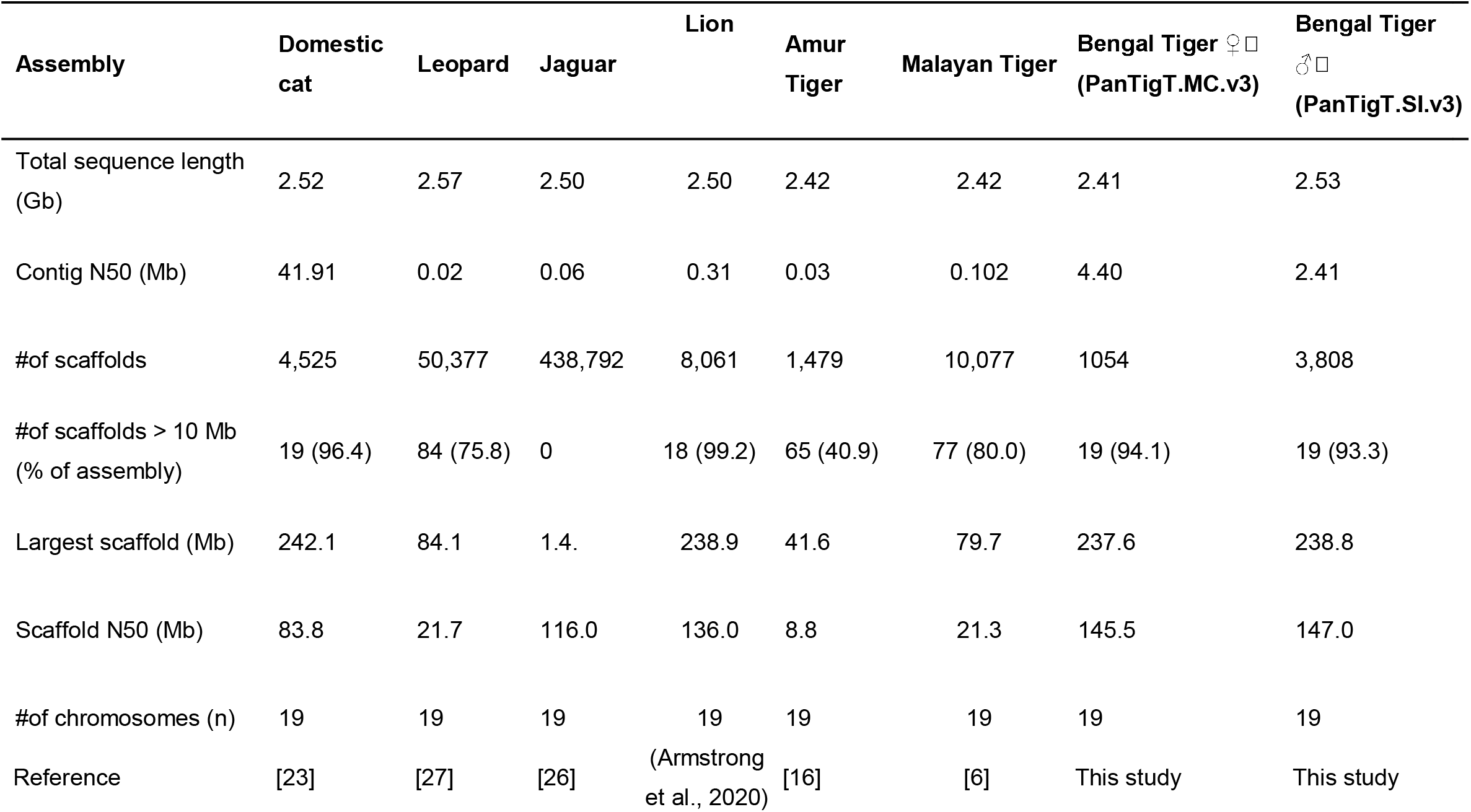
Comparison of MC and SI assemblies to other Felid genomes.

### Runs of homozygosity analysis

High-quality near-chromosomal genome assemblies can greatly help in several types of population genetic analysis, especially of threatened and endangered species. Measuring inbreeding due to shared ancestry of parental gametes is one of them. Typically, this is measured by identifying runs of homozygosity (ROH) [28–30]. We aligned short-read Illumina data from four zoo-bred Individuals with known pedigree inbreeding coefficients ranging from 0.21 to 0.28 [31] to MC, SI and the published Malayan tiger genomes (Armstrong et al. 2021) to assess the effectiveness of these assemblies in detecting ROH (**Materials and methods**). We observed on an average 2,895, 2,583 and 2,176 ROH regions that were >100 kb in length in the zoo-bred individuals when using the MC, SI and Malayan tiger genome assemblies, respectively (**Supplementary Tables 1c-e**). Further, we estimated cumulative inbreeding in individuals due to common ancestry dating back to 455, 45 and 4 generations ago (**Materials and methods**). We also observed that all three genomes performed comparably when estimating shorter ROH that represent older ancestry [7] (**Figure 1e**) while the MC and SI assemblies performed better at detection of ROH longer than 1 Mb that represents recent inbreeding, compared to the Malayan tiger genome (n= 177 for MC, n= 106 for SI vs n= 94 for Malayan tiger on average) (**Figure 1f**). All estimates of recent inbreeding from ROH longer than 1Mb correlated with the pedigree inbreeding coefficients (r^2^ = 0.97) of the individuals, thus demonstrating the need for near-chromosomal genome assemblies in conservation genomics. We conclude that near-chromosomal level genome assemblies are important for detecting long stretches of ROH that signify recent inbreeding [7] and generally host deleterious alleles.

### Genome Annotation

Our genome assemblies, PanTigT.MC.v3 and PanTigT.SI.v3, both contained 19 scaffolds >10 Mb corresponding to the 19 chromosomes (**Figure. 1a-b**). These scaffolds accounted for ~93% of the genome. The average DNA base (GC) content of the assemblies was about 41%. Analysis of the repeat content revealed that ~36% of the genome was repetitive (~870□Mb; **Figure 1a-b** and **Supplementary Table 1f** and **Supplementary Figure 2**) with long interspersed nuclear elements (LINEs) being the dominant family of repeats (**Supplementary Table 1f**).

We also annotated the genomes for noncoding RNA elements and created a database of micro RNAs (miRNAs; n=299 in MC and 301 in SI), small nucleolar RNAs (SnoRNA; n=365 in MC and 371 in SI), tRNAs (n=548 and 632 in MC and SI) and other noncoding RNA elements (**Supplementary Table 1g**). We next searched for predicted miRNAs targets using miRanda [32]. Several miRNAs were predicted to target genes involved in critical biological processes including angiogenesis (n=132 target genes; GO:0001525 and GO:0045765-0045766), brain (n=78; GO:0007420, 1990403), bone (n=73; GO:0060348, 0048539, 0030282, 0060349, 0030500) and eye (n=63; GO:0002088, 0043010, 0001654, 0048593) and heart development (n = 43; GO:0035904, 0060347, 0007507, 0007512, 0060914, 0060973) (**Supplementary Tables 1h-i**). Functional validation studies involving the predicted miRNAs [33, 34] shall lead to a better understanding of the post-transcriptional gene expression regulation in tigers and big cats during pre- and post-development.

Next, we used the MAKER pipeline [35, 36] to annotate the genomes using protein homology and RNA-seq expression data. We predicted 19,931 genes that correspond to 22,718 transcripts in the PanTigT.MC.v3 genome assembly. In the male SI individual PanTigT.SI.v3, we detected 21,126 protein-coding genes that mapped to 24,074 transcripts (**Figure 2a and Supplementary Table 1j**). Overall, the annotation yielded 26,068 unique protein-coding between the MC and SI genomes. About 95% of all annotated genes were located on the 19□largest scaffolds corresponding to the numbered chromosomes in the genomes. Using a previously developed annotation pipeline [11], we functionally classified the protein coding genes (**Supplementary Table 1j**). A total of 19,640 (86%) and 20,412 (85%) predicted proteins contained a canonical start and stop codon in the PanTigT.MC.v3 and PanTigT.SI.v3 assemblies. We found that ~99% of these genes had a corresponding ortholog in either the Human Gene Nomenclature Committee database, NCBI’s non-redundant database or the TrEMBL (https://www.ebi.ac.uk/uniprot) database (**Supplementary Tables 1k-n**). High-level gene function classification of the identified proteins revealed 179 solute carriers as the most abundant type of proteins followed by 78 zinc finger proteins (**Supplementary Figure 3** and **Supplementary Tables 1l and 1n**). Additionally, all 44 protein-coding genes identified in the Y-linked scaffolds were conserved in other Felidae genomes (43/44 genes had >90% identity) (**Supplementary Table 1o**).

**Figure 2:**
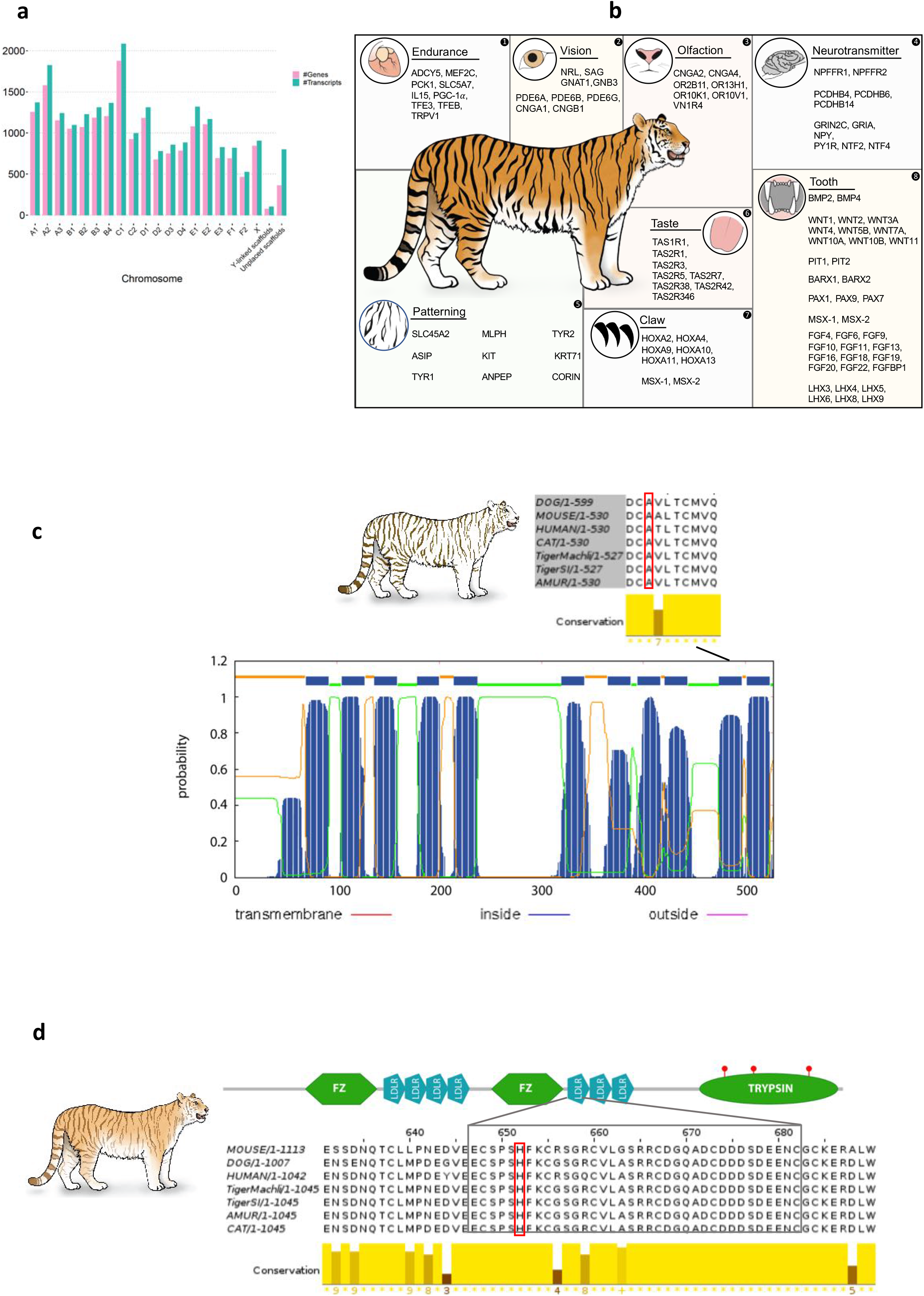
Genome Annotation and functional genes. (**a**), Bar plot of the number of predicted genes and corresponding transcripts observed in MC genome assembly and putative Y-linked scaffolds in SI genome assembly. (**b**) Putative genes involved in various key biological pathways important for apex predatory traits. (**c**) Complete *Slc45a* gene structure including the predicted transmembrane (TM) domains. Multiple sequence alignment of Scl45a c-terminal region showing the A477>V mutation observed in white tiger (**d**) Schematic diagram of full-length *Corin* gene. Multi-species alignment of the Corin LDL receptor domain depicting the known H>Y mutation associated with golden tabby pelage pattern (*Cell Res. 27(7):954–57*).

A tiger typically consumes at least one deer-sized animal each week for survival [37]. Its metabolism, sensory functions and other adaptations are key to it evolutionary success as an apex predator. Using our high-contiguity genomes, we annotated several full-length amino-acid transport (n=7; GO: 0006865) and protein, cholesterol and fatty acid metabolism-related genes (n=21; GO: 0008203, 0019538, 0006631) that likely are crucial for the successful adaptation to its carnivorous diet (**Supplementary Tables 1k and 1m**) [27]. Further, we identified multiple genes involved in G-protein coupled signaling (n=270) and olfactory receptor activity (n=107) that likely have an important role in sensory functions such as smell, vision and hearing, nervous system development, mate selection and hunting [38].

The evolutionary versatility of teeth is an important factor contributing to the success of carnivores. Diet-dependent changes in complexity of dental patterns, tooth morphology, number, function and diet are interlinked [39]. One unique morphological feature that has evolved independently numerous times within the theriodont lineage that includes mammalia, is the saber-tooth morphology of permanent upper canines [40]. Tigers have the largest upper canines of all the big cats and is key for its apex predatory status. We identified homologs of gene families known to be involved in tooth development including the *Wnt* family members and homeobox genes (*Hox*) [41] *Msx-1, Msx-2,[42], Dlx-1, Dlx-2* and *Barx-1*. We also identified *Pax9* known to be important for molar development. Additionally, our analysis identified several *Fibroblast growth factor (Fgf*), and *LIM homeodomain transcription factor (Lhx*) family members [43] involved in regulating enamel and dentin levels (**Figure 2b**; **Supplementary Tables 1k and 1m**). Mouse genetic analysis showed that the *fibroblast growth factor 10 (Fgf10*) to be a critical factor in determining the length of the teeth [44]. In addition to *Fgf genes*, we also identified *Matrix metalloproteinase-20 (Mmp-20*) that has been shown to be involved in tooth development (**Figure 2b**; **Supplementary Tables 1k and 1m**). [45, 46].

The retractable sharp tiger claws are an important tiger adaptation critical for predation. Claws are a variation in the distal limb integumentary appendages of certain mammals, including most carnivorous animals. Several epithelial-mesenchymal signaling molecules involved in patterning ectodermal derivatives such as teeth, hair, and feathers are also involved in patterning distal epidermal appendages such as claws and nails [47]. Mutations in several *Hox family member* genes have previously been shown to affect claw/nail and limb development in mice and lead to disorders such as brachydactyly in humans [1, 48–51]. In the tiger genome besides the several homeobox genes including *Homeobox* family genes (*Hox), Msx-1* and *Msx-2*, we identified bone morphogenic protein (*Bmp*) family members known to be involved in early distal limb development (e.g., *Bmp4*, posterior *Hox* genes) and induction of the claw growth, and/or proliferation (**Figure 2b**; **Supplementary Tables 1k and 1m**).

It is estimated that about two-thirds of all extant carnivorous mammals are mainly nocturnal [52]. Nocturnality is characterized by an expansion in the number of genes responsible for dim/low-light vision. The retina is a light-sensitive layer of eye tissue and consists of six types of neurons which include light-sensitive photoreceptor cells (rods and cones). Rod cells are responsible for discerning shapes and are highly sensitive to low light while cone cells confer bright light sensitivity as well as color vision. Tigers have more rods in their eyes than cones [53]. We identified several genes critical for low-light vision, including Neural retina leucine zipper (*Nrl*), S-antigen visual arrestin (*Sag*), G Protein Subunit Alpha Transducin 1 (*Gnat1*), G Protein Subunit Beta 3 (*Gnb3*), Phosphodiesterase genes *Pde6a, Pde6b and Pd36g*, and Cyclic Nucleotide Gated Channel Subunit Alpha 1 (*Cnga1*) and Beta 1 (*Cngb1*) (**Figure 2b**; **Supplementary Tables 1k and 1m**). Knockout mutation of *Nrl* leads to the loss of rod cells [54] while mutations in *Gnb3, Cnga1* and *Cngb1* have been shown to lead to night blindness in humans and retinal degeneration in chickens [55–58].

While the tiger’s sense of smell, though not critical for hunting, is used for communication between animals particularly, for marking territory, courtship and reproduction. The vomeronasal organ, also called the Jacobson’s organ, is primarily responsible for pheromone detection in tigers [59]. We identified twenty one vomeronasal type-1 receptor genes (V1R), of which twelve were incomplete with truncated sequences, indicating potential pseudogenization, consistent with findings in other carnivorous species [60]. In contrast to the ~400 olfactory receptor genes found in humans [61], we identified 69 olfactory receptor (OR) genes based on homology to other OR genes (**Figure 2b**; **Supplementary Tables 1k and 1m**) [48].

Mammalian taste is mainly mediated by receptor cells organized in taste buds on the tongue [62]. It has been established that the tiger can taste salt, bitter and acidic flavors and to a lesser degree sweetness [63, 64]. We identified a fully intact copy of *Tas1r1* (type 1 taste receptor), responsible for sweet taste perception, and two Tas2r (type 2 taste receptor) genes involved in bitter taste perception [65] (**Figure 2b**; **Supplementary Tables 1k and 1m**). Notably, we did not identify a full-length homolog of *Tas1r2*, a sweet taste receptor gene, consistent with observations in other carnivorous mammals [64, 66].

Tigers are the only striped cat of the genus *Panthera* and are most recognizable for their pattern of black vertical stripes on reddish-orange fur. Our high-quality assembly and annotation identified several genes associated with morphometric variation in domestic cats such as pigmentation, coat patterns and other phenotypes. Included among these were homologs of the atrial natriuretic peptide-converting enzyme (*Corin*), membrane-associated transporter protein (*Slc45a2), Agouti-signaling protein (Asip), Tyrosinase-related protein 1 (Tyrp1), Tyrosinase-related protein 2* (*Tyrp2*), *Tyrosinase* (*Tyr*), *Melanophilin* (*Mlph*), *Tyrosine-Protein Kinase Kit* (*Kit*), *Keratin 71 (Krt71*) and *Aminopeptidase N (Laeverin; Anpep*). Mutations in these genes can lead to a variety of phenotypes including variation in coat and feet color, stripe patterns and the color of stripes, tail shapes (**Figure 2c-d**) [1, 67–71]. Of note, our high-quality genome enabled accurate annotation of long genes with their complete intron-exon structure including that of *Slc45a2*, a ~33kb gene consisting of 7 exons. Using a combination of transcriptome (RNA-seq) and genome data, we were also able to annotate a ~249 kb long full-length copy of *Corin* made of 22 exons. These fully annotated genes will serve as a high-quality resource for understanding coat patterning genes in tigers and other animals.

### Phylogeny and positive selection analysis

To compare the annotations of the genomes from this study to other big cat species including the Amur tiger, we constructed a phylogeny using annotated protein sequences from 12 selected species, namely, dog, Amur tiger, Bengal tigers (this study), lion, leopard, jaguar, clouded leopard, domestic cat, Canada lynx, cougar, horse, and rabbit. The proteins from these species clustered into 26,169 orthogroups (representing 97.2% of the input protein sequences) of which 4,576 were single copy orthologs. The remaining protein sequences did not cluster (854 proteins (3.5%) in SI and 489 genes (2.2%) in MC, respectively). Next, we used the 4,576 single-copy orthologs to test for signals of positive selection in big cats using site models in PAML [72]. Using Likelihood Ratio Tests (LRT) for models M1-M2 and M7-M8, we identified 1,484 single copy orthologs to be positively selected (p-value <0.05) (**Supplementary Tables 1p-r**). Included in this list of positively selected genes, were genes involved in muscle development *Atp1b4*, angiogenesis (*Srpx2*), fatty acid and cholesterol metabolism (*Slc4a9, Apod*), mitochondrial respiration (*Cq10a*) and heart development/endurance-related genes (e.g., *Nkx25*), indicating that these genes evolved at a faster rate, consistent with the physiological needs of big cats (**Figure 3b** and **Supplementary Tables 1p-r**) [16, 26, 27, 38]. Further, pathway enrichment analysis of the positively selected genes confirmed enrichment of several metabolism-related pathways including cholesterol metabolism (GO:0008203) and fatty acid metabolism (GO:0006631) that are critical for obligate carnivores (**Figure 3c**).

**Figure 3:**
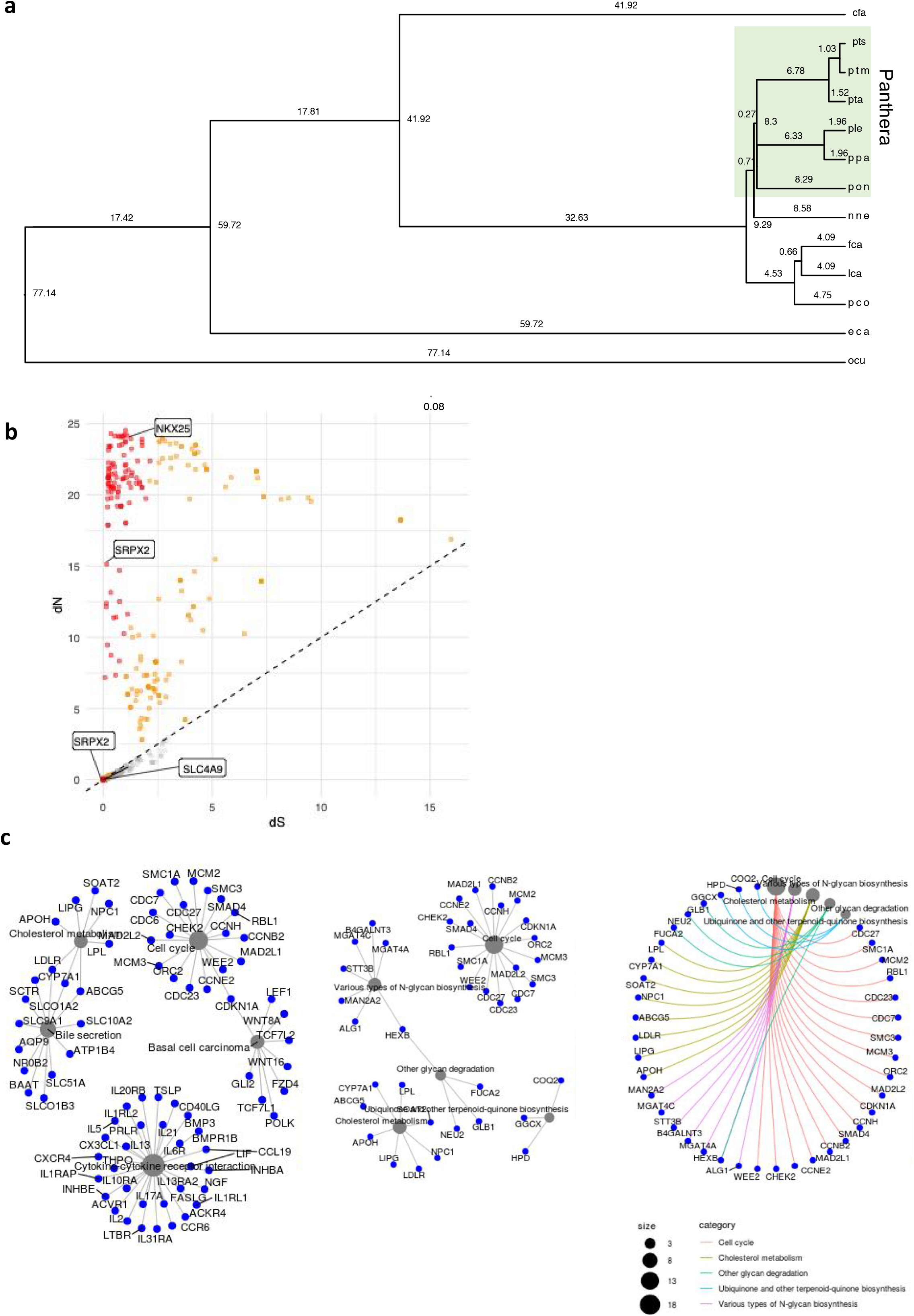
Phylogeny and positive selection analysis. (**a**) Phylogeny tree constructed using single copy orthologs from selected Panthera and outgroup species. **(b)** Scatter plot of evolutionary rates of single copy key pattern and pelage-related gene orthologs identified in the Bengal tiger genome and **(c)** Pathway analysis of single copy orthologs within the Panthera lineage showing gene network interaction models. cfa: *Canis familiaris* (dog), pts: south Indian Bengal tiger (male), ptm: Machali Bengal tigress (female), pta: *Panthera tigris altaica* (Amur tiger), ple: *Panthera leo* (lion), ppa: *Panthera pardus* (leopard), pon: *Panthera onca* (jaguar), nne: *Neofelis nebulosa* (clouded leopard), fca: *Felis catus* (domestic cat), lca: *Lynx canadensis* (Canada lynx), pco: *Puma concolor* (cougar), eca: *Equus caballus* (domesticated horse), ocu: *Oryctolagus cuniculus* (rabbit)

## Discussion

In this study, we have generated de novo genome assemblies and protein-coding annotations for a female and a male Bengal tiger. The resulting assemblies were 17x more contiguous than the published Amur tiger genome, ~7x more contiguous than the Malayan tiger genome, and 1.7x more contiguous than the domestic cat genome with a scaffold N50 of over 140□Mb, making them the most contiguous near-chromosomal wild felid genomes assembled to date (**Table 3**).

We provide here a comprehensive annotation of 26,068 protein-coding genes from the tiger genomes. Additionally, we identified over 3,000 non-coding genes including for the first time a genome-wide analysis of micro-RNAs and their putative target genes in a tiger genome. Functional assignment identified genes and signaling pathways involved in endurance, neurotransmission, teeth, vision, claw and skin patterning, all of which are important for the tiger’s adaption as an apex predator. We have performed extensive annotation of genes involved in variations in coat color, stripe patterns and other morphometric variations.

Our highly contiguous genome along with the annotations is an important resource for studies on the genetics of Felidae and in general carnivoran development and evolution. For example, it is now feasible to establish homology between tooth components between different big cat species by using the annotation generated in this study. Our annotated gene sets for the Bengal tiger genomes provides a high-quality reference that can be used to support resequencing efforts aimed at studying tiger populations. The genome assemblies presented here demonstrate their utility in population and conservation genetic studies of tigers. They increase the accuracy of the estimates of important population genetic parameters like inbreeding for assessing the threats to a population. Our reference genome along with annotation-guided quantitative studies on wild and inbred, zoo-bred individuals will help in understanding the possible phenotypic effects of inbreeding depression through assessment of ROH (Leroy 2014). The high-quality genomes present in this study will be important resources that will enable analysis of sequencing data obtained from DNA present in non-invasive scat samples obtained from the wild. Such analysis will be important for understanding and monitoring tiger population genetic diversity in the wild and support informed conservation efforts.

## Methods

### Samples and nucleic acid preps

Bengal tiger blood samples used in the study are reported in **Supplementary Table 1a**). The SI blood samples were collected from an animal in Wayanad Wildlife Sanctuary, Kerala, India under the order of the Chief Wildlife Warden of Kerala on April 12^th^, 2015. MC and ST3 samples used in the study were previously described in [7]. Genomic DNA from MC and SI was extracted from whole blood using the MagAttract HMW DNA extraction kit (Qiagen). Sample quality was checked using Qubit 3.0, Nanodrop 8000, Agilent TapeStation 4200 and pulse field gel electrophoresis (Sage Science). Total RNA was isolated from the ST3 blood using Trizol (ThermoFisher). A total of 500ng of RNA was used for library preparation.

### Sequencing

Sequencing libraries for PacBio, Oxford Nanopore (ONT) and Illumina sequencing were constructed as per manufacturer’s instructions. Considering the genome size of *Panthera tigris* at ~2.4 Gb, we generated ~79.6 Gb of Pacbio long-read data (~33x; N50 = 10.3 Kb) and ~18.28 Gb of ONT long-read data (~7.5x, N50 = 8.14 kb) for the female tiger (MC). In addition, short-read Illumina sequencing data 117 Gb, (50x; 250 bp library), 35 Gb (15x; 300 bp library 35 Gb (15x; 500 bp library), and 35 Gb (15x; 800 bp library) was also generated. For the male tiger (SI), a total of 160.3 Gb (64x) ONT data, and ~60 Gb (25x) of Illumina short-read data was generated (**Supplementary Table 1b**). In addition, blood from an offspring of Machali, ST3 was also obtained.

RNA-seq short read (Illumina) data was obtained by sequencing PolyA RNA-sequencing (RNA-seq) libraries prepared from ST3 blood using the Illumina TruSeq stranded messenger RNA kit (**Supplementary Table 1b**).

### Chicago library preparation and sequencing

Chicago library was prepared as described previously [20]. Briefly, for each library, ~500ng of HMW gDNA (mean fragment length of 75 kb) was reconstituted into chromatin *in vitro* and fixed with formaldehyde. Fixed chromatin was digested with DpnII, the 5’ overhangs filled in with biotinylated nucleotides, and then free blunt ends were ligated. After ligation, crosslinks were reversed and the DNA purified from protein. Purified DNA was treated to remove biotin that was not internal to ligated fragments. The DNA was then sheared to ~350 bp mean fragment size and sequencing libraries were generated using NEBNext Ultra enzymes and Illumina-compatible adapters. Biotin-containing fragments were isolated using streptavidin beads before PCR enrichment of each library. The libraries were sequenced on an Illumina platform. A total of 155 million read pairs (2 x 150 bp) was generated for this library providing ~20x physical coverage of the genome (1-50 kb pairs) (**Supplementary Table 1b**).

### Bionano optical mapping data generation

High-molecular weight genomic DNA extracted from whole blood from SI was embedded in a thin agarose layer and was labeled and counterstained following the Direct Label and Stain (DLS) Reagent Kit protocol (Bionano Genomics). The labeled sample was then loaded onto a Saphyr chip and run on the Saphyr imaging instrument (BioNano Genomics). A total of 473.7 Gb (~190x) optical map data was generated. *De novo* genome assembly using the Bionano Access software for the DLS optical map data produced a 2.63 Gb assembly (PanTigT_BNG) consisting of 182 scaffolds (N50 =145.93 Mb) (**Supplementary Table 1b**).

### Hi-C library preparation and sequencing

Hi-C library (Dovetail Genomics), was prepared for the two samples (SI and ST3) as described previously [19]. Briefly, chromatin was fixed in place with formaldehyde in the nucleus and then extracted. Fixed chromatin was digested with DpnII, the 5’ overhangs filled in with biotinylated nucleotides, and then free blunt ends were ligated. After ligation, crosslinks were reversed and the DNA purified from protein. Purified DNA was treated to remove biotin that was not internal to ligated fragments. The DNA was then sheared to ~350 bp mean fragment size and sequencing libraries were generated using NEBNext Ultra enzymes and Illumina-compatible adapters and sequenced (**Supplementary Table 1b**).

### Genome assembly

We used Canu (v1.7.1; *corMinCoverage* = 0, *corMhapSensitivity = high and correctedErrorRate=0.12* (MC)/0.15 (SI)) to generate an initial draft assembly for both the study animals. The primary MC draft assembly (PanTigT.MC.v1) consisted of 2.4 Gb spanning 2,766 contigs (contig N50 = 4.40 Mb) (**Table 1**) and the SI draft assembly consisted of 845 contigs totaling 2.38 Gb (contig N50 = 29.03 Mb). Next, both assemblies were polished to correct for base call errors using raw PacBio data (MC only) and Illumina data (MC and SI). Briefly, PacBio reads were aligned to the PanTigT.MC.v1 using *pbalign* (pbalign version: 0.3.1) resulting alignment bam file was used as input for polishing using *Arrow* (https://github.com/pacificbiosciences/genomicconsensus/). A total of two rounds of polishing were done using *Arrow*. Illumina short-read libraries generated for MC and SI (**Supplementary Table 1b**) were used for error correction and conflict resolution using Pilon (v1.22) with the following parameters (*--changes --diploid --fix all*) to produce polished assemblies PanTig.T.MC.v1 and PanTigT.SI.v1 (**Table 1**).

Next, PanTigT.MC.v1 was used as the input for scaffolding with Hi-C sequence data using HiRise resulting in an assembly PanTigT.MC.v2 containing 1,052 scaffolds (scaffold N50= 145.7 Mb; **Supplementary Figure 1a**). Similarly, PanTigT.SI.v1 was used as the input for scaffolding with Chicago sequence data using HiRise. This resulted in an assembly PanTigT.SI.v2 containing 6,889 scaffolds (scaffold N50 = 2.3 Mb). PanTigT.SI.v2 was then scaffolded with Bionano optical map data to yield PanTigT.SI.v3 assembly comprising of 3,886 scaffolds (scaffold N50 = 141.3 Mb). Interestingly, incorporation of Hi-C data did not improve the contiguity of PanTigT.SI.v3 (**Supplementary Figure 1b**) and did not result in any breaks or joins to the assembly. We also assessed if the order of integration of the different sequencing data types would affect assembly quality. While scaffolding PanTigT.SI.v1 with Hi-C data alone (no Bionano and Chicago) resulted in an assembly with fewer scaffolds (n=692) but lower N50 (125.3 Mb), scaffolding of PanTigT.SI.v1 with only Chicago and Hi-C data resulted in an assembly with 6,461 scaffolds and lower scaffold N50 of 48.2 Mb. Thus, we retained the Chicago-scaffolded assembly (PanTigT.SI.v2) for downstream analysis including chromosome assignment. Using the domestic cat reference assembly (Felis_catus_9.0), we merged scaffolds into chromosomes, corrected orientation errors and assigned chromosomes based on synteny using RaGOO (https://github.com/malonge/RaGOO) resulting in the final SI assembly PanTigT.SI.v3 and PanTigT.MC.v3.

Next, error-corrected reads obtained from long-read data were used in conjunction with RAILS (v1.4.1) (https://github.com/bcgsc/RAILS) to fill gaps in PanTigT.MC.v3 and PanTigT.SI.v3 using Cobbler with the following parameters: -d 1000 -i 0.95 [73]. A total of 21 out of 1700 gaps (1.24%) were filled in PanTigT.MC.v3 with the largest gap closed being ~10,025 bp while no significant improvement was observed for PanTigT.SI.v3. A final round of polishing with Racon (v1.3.3) [74] was done on PanTigT.MC.v3. To assess genome similairy, we mapped the two genome assemblies with minimap2 and called variants with paftools.js. Briefly, using the MC genome as the reference, we aligned SI genome and detected 2,196,239 SNPs per 2,087,495,282 aligned bases. This translates to ~1.05 SNPs/kb of the SI genome.

### Y-chromosome-linked scaffold identification

To identify the male Y-linked scaffolds, we separately aligned Illumina reads obtained from the female (MC) and the male animal (SI) to PanTigT.SI.v3 assembly using BWA [75] allowing for two mismatches and one indel. Scaffolds with less than 80% alignment coverage were excluded from further analysis. Then, single-base depths were calculated using SAMtools [76] following which coverage and mean depth for each scaffold was calculated. Using the average coverage across a scaffold using either the male or female reads, we identified 3 Y-linked scaffolds Sc4gRnz_35;HRSCAF=35_pilon, Sc4gRnz_53;HRSCAF=53_pilon and Sc4gRnz_8;HRSCAF=8_pilon.

### Chromosome painting

Chromosome painting of SI or MC chromosomes with the domestic cat chromosomes was performed using the SatsumaSynteny2 script with default parameters [77].

### Runs of Homozygosity (ROH) analysis

ROH were identified as described in Armstrong et al. (2021). We aligned reads from short-read sequencing (**Supplementary Table 1b**) from four zoo-bred tigers with known inbreeding coefficients [31] to MC, SI and Malayan tiger genome assembly (Armstrong et al. 2021). After sequence alignment, variant calling and filtering we used BCFtools/ROH at default settings to determine the allele frequencies. We categorized ROH into three categories of stretches longer than 100Kb, 1Mb and 10Mb.

### Repeat element identification

We identified the repetitive elements in the genome by combining both homology-based and de novo predictions. Briefly, we used RepeatModeler (v.1.0.11) (http://www.repeatmasker.org/RepeatModeler.html) to construct the species-specific repeat sequence libraries for both SI and MC genome assemblies, and then used these as a query to identify repetitive elements using RepeatMasker.

### Noncoding RNA Annotation

tRNA-encoding genes were predicted by tRNAscan-SE (v1.3.1) [78] with default parameters. Ribosomal RNA (rRNA) fragments were predicted by aligning to mammalian rRNA sequences by using BLASTN (v 2.2.26) with an E-value cutoff of 1e-10. MicroRNA (miRNA) and small nuclear RNA (snRNA) genes were annotated using INFERNAL (v1.1.3) [79] by searching against the Rfam database (release 13.0) [80]. Whole-genome target gene prediction for miRNAs was performed using miRanda (v3.3a) [32] using default parameters. Gene ontology terms for the target genes were obtained from the functional annotation for the genomes.

### Genome Annotation

Gene prediction was performed on the assemblies PanTigT.SI.v3 and PanTigT.MC.v3 using the genome annotation tool, MAKER2 (v2.31.10) [35] in an iterative process. First, *ab initio* gene prediction was performed by the programs SNAP (v2006-07-28) [81] and Augustus (v3.2.3) [82] using publicly available transcriptome data from the Siberian tiger (SRA091968), a de novo assembled transcriptome from ST3 blood RNA-seq data using Trinity [83] and a combined protein database consisting of proteins from the UniProt/Swiss-Prot and NCBI non-redundant databases of reviewed proteins. A total of three iterative runs of MAKER was used to refine the gene models and produce the final gene set with an annotation edit distance (AED) cutoff of 0.6. Genome annotation quality was assessed by BUSCO analysis using the conserved core set of mammalian genes (79.3% and 76.9% BUSCO completeness score for MC and SI assemblies, respectively). Next, using the set of complete predicted protein sequences, we performed functional annotation as described previously [11] (**Supplementary Table 1g**).

To identify gene families specific to tooth development, claw, taste, neurotransmitter, olfaction, and coat color/pattern, we performed a literature-based search [1, 26, 48, 49] to identify genes potentially involved in these developmental pathways. Completeness of gene models for each trait-specific gene where present (predicted by MAKER) was confirmed by aligning each gene against its homolog using NCBI BLAST. Genes that were partial or incompletely annotated were manually curated. Briefly, for each incomplete gene, the corresponding homolog obtained from UniProt was aligned to the genome (SI/MC) using the ‘est2genome’ parameter in the Exonerate program with default parameters. The aligned genome coordinates of trait-specific genes were then checked against the annotation file to validate the gene prediction.

### Phylogeny and positive selection analysis

Proteomes from 12 selected species, namely, dog, Bengal tiger (this study), Amur tiger, lion, leopard, jaguar, clouded leopard, domestic cat, Canada lynx, cougar, horse, and rabbit were obtained from Ensembl Biomart (www.ensembl.org). Next, Orthofinder (v2.5.2) was used to cluster the proteins into orthogroups. Of note, the snow leopard was excluded from this analysis due to limited annotation models and lack of a reference genome. Next, single copy orthologs were extracted for evolutionary divergence estimation and positive selection analyses. The single copy orthologs were aligned with Muscle (v3.8.31) and alignments trimmed and converted to codon level alignments using trimal (v1.4rev15). For each of these alignments PAML/codeml (v.4.10) [72] was run pairwise to obtain four-fold degenerate bases which are concatenated into a super-matrix based on gene partitions. The dN/dS ratios are also obtained in this process. The resulting supermatrix was used to generate a species tree using iqtree2 (v2.0.3) with GTR+F+I+G4 models along with ultrabootstrap (1000 iterations). Further, PAML/mcmctree was used to estimate evolutionary divergence among the species considered with 95% confidence intervals with a burn-in of 2,000 and sampling 20,000. The evolutionary divergence calibrations between the following species pairs were extracted from timetree.org: *Felis catus* and *Lynx canadensis* (7.9 - 13.1 MYA), *Puma concolor* and *Felis catus* (8.8 - 13.9 MYA), *Panthera leo* and *Panthera pardus* (2.6 - 4.74 MYA), *Neofelis nebulosa* and *Panthera onca* (7.4 - 14 MYA).

### Site model tests

The single copy orthologs were fit to the following codon substitution models (M0, M1, M2, M7, M8) of PAML/codeml [72]. Likelihood Ratio Test (LRT) between M1 vs M2 and M8 vs M7 pairs were used to determine positive selection at Chi-square pvalue <= 0.05. Multiple testing using Benjamini-Hodgberg (BH) method was then performed to determine FDR (False Discovery Rate) for the p-values obtained above to determine significantly positively selected genes among the species. Bayesian Empirical Bayes (BEB) [84] results were used to identify the sites of positive selection at >=95% and >=99% confidence intervals.

## Software URLs

Canu - https://github.com/marbl/canu

RAILS - https://github.com/bcgsc/RAILS

RaGOO - https://github.com/malonge/RaGOO

PAML – http://abacus.gene.ucl.ac.uk/software/paml.html

Orthofinder - https://github.com/davidemms/OrthoFinder

Trimal – http://trimal.cgenomics.org/

Iqtree - http://www.iqtree.org/

Muscle - https://www.ebi.ac.uk/Tools/msa/muscle/

Vcftools - http://vcftools.sourceforge.net/

VEP – https://www.ensembl.org/vep

MAKER - http://www.yandell-lab.org/software/maker.html

BUSCO - https://busco.ezlab.org/

Exonerate - https://www.ebi.ac.uk/about/vertebrate-genomics/software/exonerate

Trinity - https://github.com/trinityrnaseq/trinityrnaseq

Augustus - http://bioinf.uni-greifswald.de/augustus/

SNAP - https://github.com/KorfLab/SNAP

Repeatmasker - https://www.repeatmasker.org/

RepeatModeler - https://www.repeatmasker.org/

D-Genies - http://dgenies.toulouse.inra.fr/

Satsuma - https://github.com/bioinfologics/satsuma2

Pilon - https://github.com/broadinstitute/pilon

miRanda - http://cbio.mskcc.org/miRNA2003/miranda.html

INFERNAL - http://eddylab.org/infernal

PLINK - http://zzz.bwh.harvard.edu/plink/

BWA - https://github.com/lh3/bwa

Samtools - http://www.htslib.org/

Sentieon - https://github.com/Sentieon

Racon - https://github.com/isovic/racon

## Data Accessibility

Raw sequencing data (DNA and RNA) and genome assemblies can be accessed at NCBI under BioProject accession numbers PRJNA732096 and PRJNA796358.

## Acknowledgements

We would like to acknowledge Dovetail genomics for sponsoring the HiC kits and for bioinformatics analysis using the HiRise platform. We would like to thank Subhadeep Bhattacharya and Kaushalkumar Patel for help with sample collection. We also thank the Rajasthan and Kerala Forest Department for their support.

## Author contributions

**H.S.**: Formal analysis, Data curation, Visualization, Writing – original draft, reviewing and editing **K.S.**: Formal analysis, Data curation, Visualization, Writing – original draft, reviewing and editing **A.K.**: Conceptualization, Formal analysis, Visualization, Investigation, Methodology, Resources, Writing – original draft, reviewing and editing **K.M.**: Formal analysis, Visualization, Writing – original draft **R.C.P.**: Formal analysis **O.K.M.**: Formal analysis, Data curation, Visualization **R.M.**: Formal analysis, Writing – original draft **M.D.D.**: Data curation **M.M.**: Investigation **B.K.**: Investigation, Resources **S.M.**: Investigation **S.P.K.**: Resources **A.Z.**: Resources **S.S.**: Conceptualization, Supervision, Project administration, Writing – original draft, reviewing and editing **U.R.**: Conceptualization, Supervision, Project administration, Writing – original draft, reviewing and editing

