## supplementary figures for "Near-chromosomal *de novo* assembly of Bengal tiger genome reveals genetic hallmarks of apex-predation"

### 03. Supplementary Figures

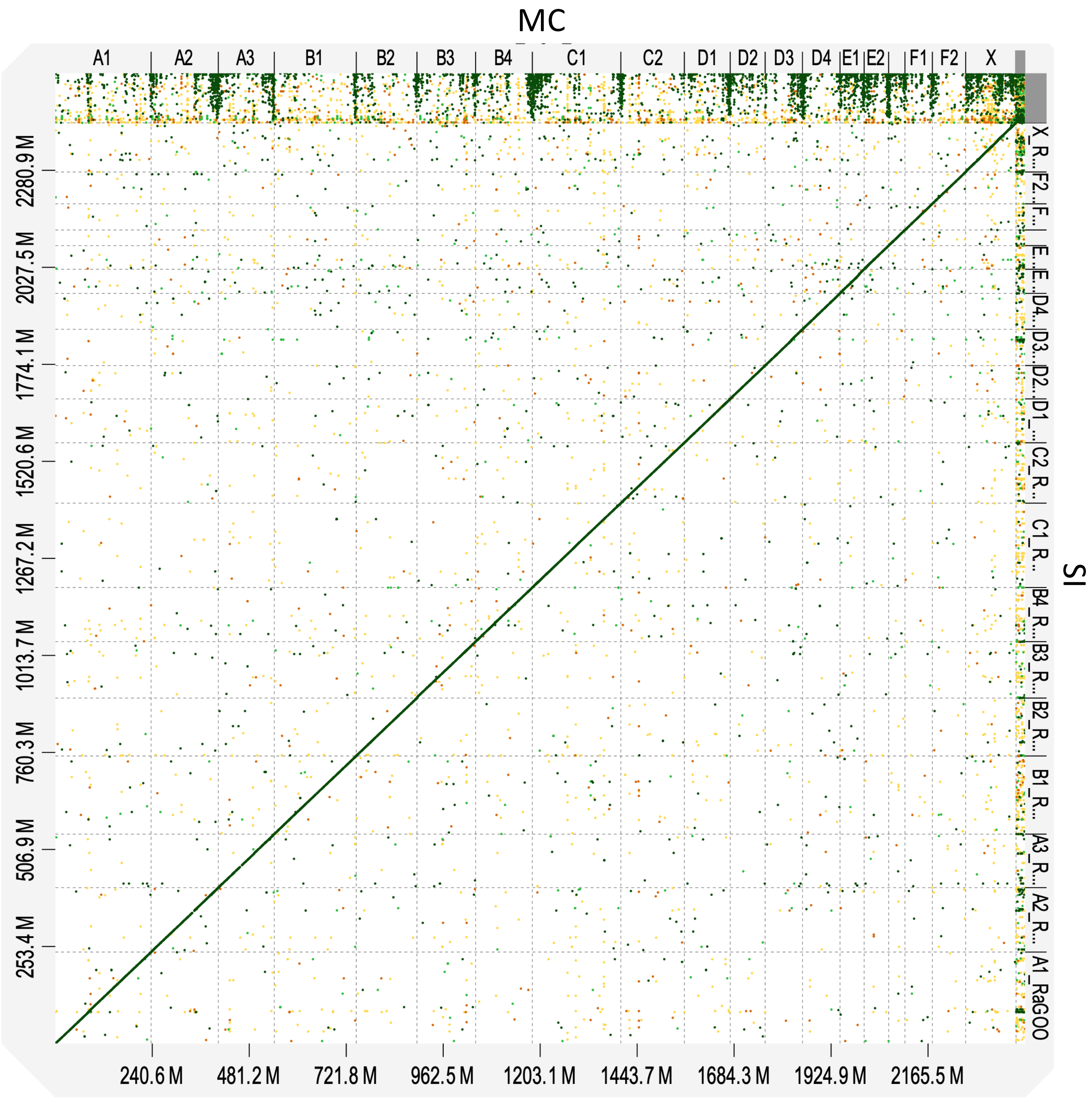


**Supplementary Figure 1:** Synteny between MC and SI tiger genome assemblies

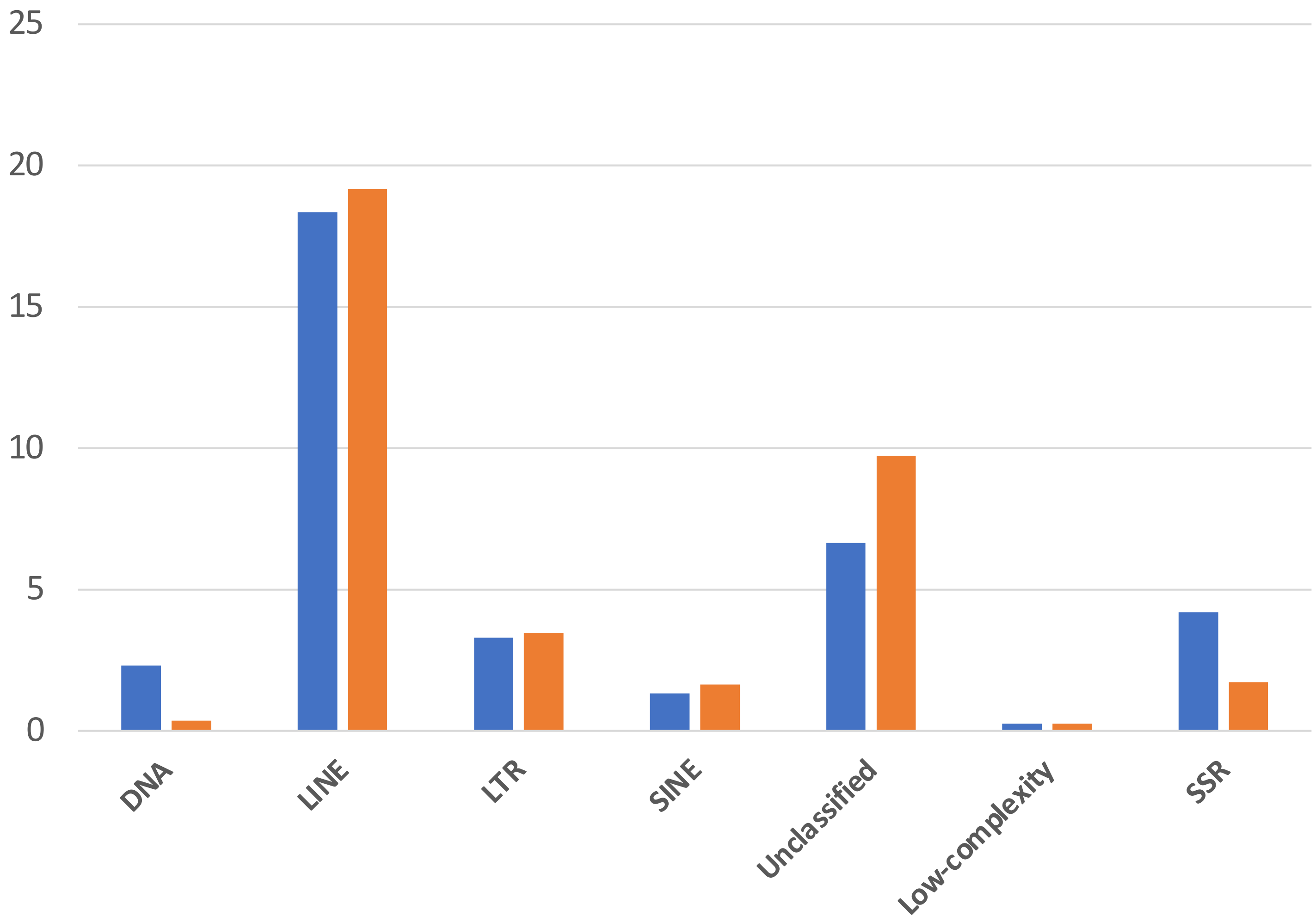


**Supplementary Figure 2.** Repeat element distribution in MC (blue bars) and SI (orange bars) genomes. LINE: long interspersed nuclear element; LTR: long terminal repeat; SINE: short interspersed nuclear element, SSR: simple short repeat

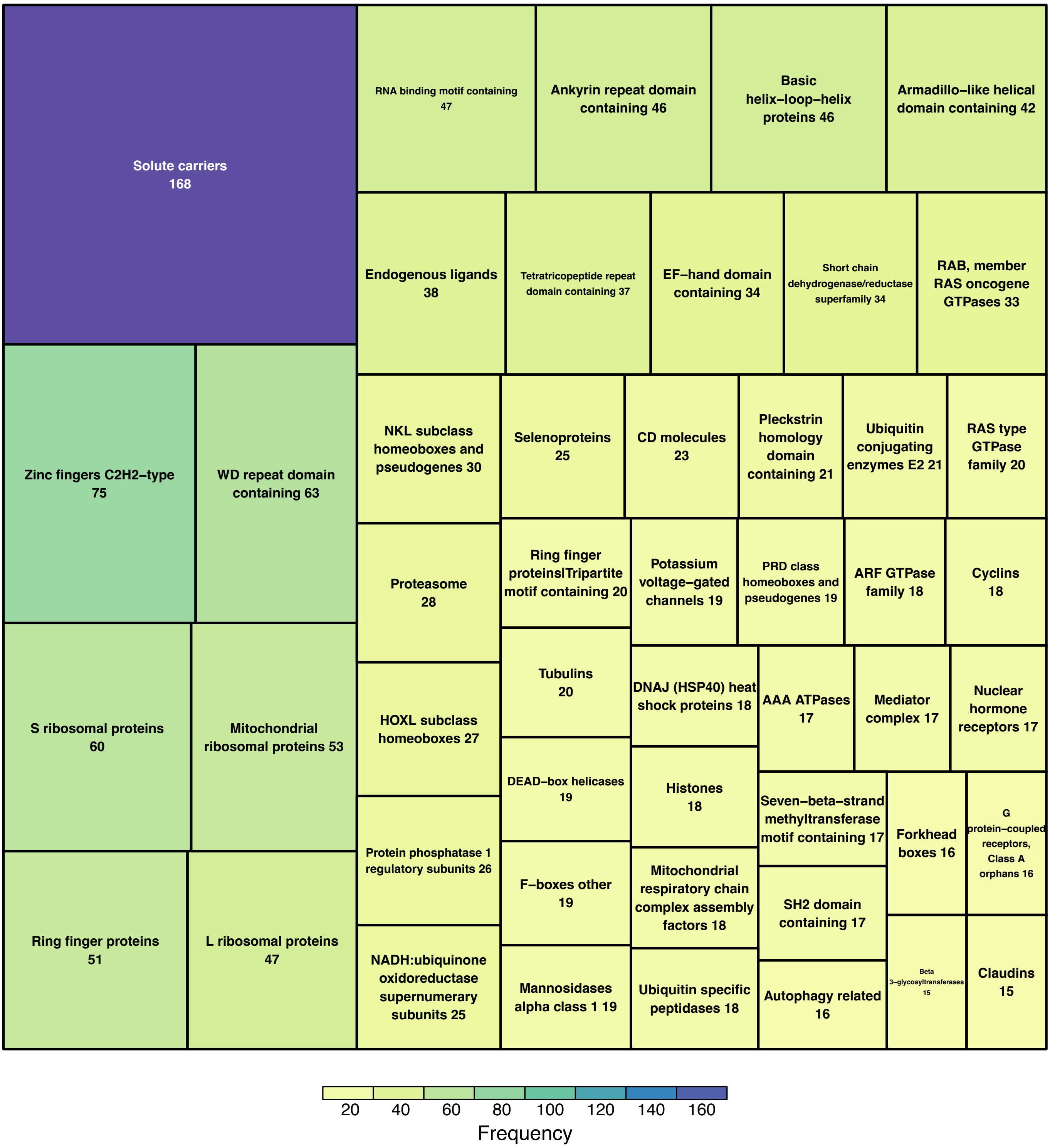


**Supplementary Figure 3.** Treemap plot of gene families identified in MC genome

a)

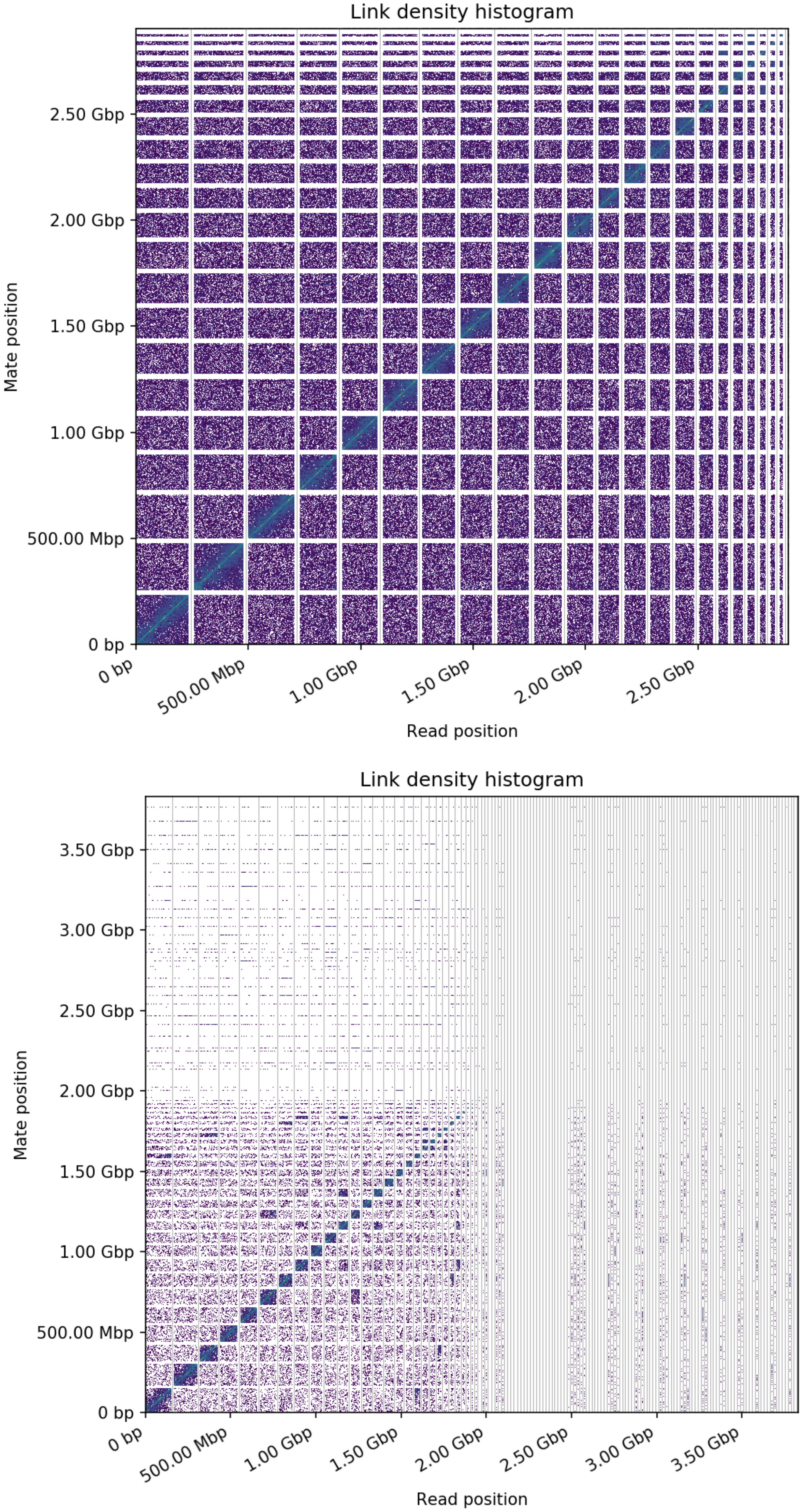


b)



**Supplementary Figure 4.** Contact maps of (a) PanTigT.MC.v2 and (b) PanTigT.SI.v3. the x and y axes give the mapping positions of the first and second read in the read pair respectively, grouped into bins. The color of each square gives the number of read pairs within that bin. White vertical and black horizontal lines have been added to show the borders between scaffolds. Scaffolds less than 1 Mb are excluded.
